# Functional differences drive the selection of NRAS mutants in melanoma

**DOI:** 10.1101/2021.01.15.426808

**Authors:** Brandon M. Murphy, Tirzah J. Weiss, Andrea M. Holderbaum, Aastha Dhakal, Marie Fort, Michael S. Bodnar, Min Chen, Craig J. Burd, Vincenzo Coppola, Christin E. Burd

**Author notes:** **CORRESPONSING AUTHOR:** Christin E. Burd, Biomedical Research Tower, Rm 918, The Ohio State University, Columbus, Ohio 43210, USA. **CONTRIBUTIONS:** B.M.M and C.E.B conceived the study and wrote the manuscript. M.C and V.C performed CRISPR-Cas9 targeting to develop the NRAS-mutant TN mouse models. B.M.M, C.E.B., T.J.W, A.M.H and M.F. generated the experimental mouse colonies, tracked tumor formation and contributed to the analysis of in vivo data. B.M.M isolated primary cells and performed in vitro signaling assays with help from A.D. and M.S.B. B.M.M and C.J.B. processed and analyzed the RNA sequencing data. All authors assisted in editing the manuscript.

## Abstract

Distinct NRAS mutants are enriched in various tumor types. Here, we generated a suite of fully congenic, conditional, *Nras* knock-in mouse models (*LSL-Nras Q61R, -K, -L, -H, -P, -Q; G12D* and *G13D, -R*) to test the hypothesis that melanocyte transformation requires functions specific to the NRAS mutants enriched in human melanoma (Q61R and Q61K). Consistent with the rarity of NRAS codon 12 and 13 mutants in human melanoma, spontaneous melanomas were rare or absent in mice expressing NRAS G12D, G13D or G13R. Mice expressing less common codon 61 alleles (Q61H, Q61P) also developed few or no tumors. NRAS Q61R, Q61K, or Q61L expression, by contrast, induced rapid melanoma onset with high penetrance. Cohorts of heterozygous mice containing one *LSL-Nras Q61R* and one *LSL-Nras Q61K, -L, -H, -P*, or *-Q* allele were generated to assess potential interactions between NRAS mutants. The ability of each *Nras* variant to substitute for an *Nras Q61R* allele was consistent with its own ability to drive spontaneous melanoma formation. However, *LSL-Nras Q61Q*/*Q61R* mice rarely developed tumors. *In vitro* experiments in mouse embryonic fibroblasts (MEFs) highlighted activation of the MAPK pathway as a defining difference between tumorigenic and non-tumorigenic NRAS mutants. Enhanced MAPK activation was associated with the promotion of BRAF-BRAF and BRAF-CRAF dimers. These results support the development of cancer preventative strategies specific to the properties of the commonly observed RAS mutants in each tumor type.

## INTRODUCTION

It is unclear why the profile of oncogenic *RAS* mutations differs between tumor types. It was once thought that differences in tumor etiology determined the preferred location (codon 12, 13 or 61) and amino acid identity of oncogenic mutations in *RAS*. However, apart from *KRAS*^*12C*^ mutations which are linked to cigarette carcinogens in lung cancer (Dogan et al. 2012), tumor type-specific mutational processes do not explain the enrichment of specific *RAS* mutations in many cancers. This trend is particularly evident in melanoma where the most common *NRAS* mutations (Q61R and Q61K) are not caused by direct damage from ultraviolet (UVB) light (Hodis et al. 2012). These observations suggest that each RAS mutant may fulfill different requirements for tumor initiation.

Emerging evidence shows that RAS mutants have distinct biochemical and tumorigenic properties. While all oncogenic RAS mutants are constitutively active, differential positioning of the switch I and II domains leads to variances in GTP binding and hydrolysis (Lu et al. 2016; Novelli et al. 2018). These structural differences can also influence effector interactions (Céspedes et al. 2006; Buhrman et al. 2007; Stolze et al. 2014; Hunter et al. 2015; Marcus and Mattos 2015) as evidenced by the positioning of switch II in KRAS^12R^, which prevents PI3Kα binding and the subsequent induction of macropinocytosis (Hobbs et al. 2020). Such mechanistic differences may also explain the tissue-specific potential of RAS mutants to initiate tumorigenesis in genetically engineered mouse models (GEMMs). For example, we have shown that endogenous levels of NRAS^61R^ or NRAS^12D^ exhibit distinct tumorigenic potential in GEMMs of melanoma and leukemia (Burd et al. 2014; Kong et al. 2016). Finally, mutation-specific functions of oncogenic RAS may influence patient outcomes as the efficacy of targeted therapies in colorectal and non-small cell lung cancer is dependent upon the underlying KRAS mutant (de Roock et al. 2010; Tejpar et al. 2012; Mao et al. 2013; Bournet et al. 2016). Therefore, understanding functional differences that drive the selection of specific RAS mutants in each cancer type may identify pharmacologically tractable targets required for tumor initiation.

Technical challenges have made it hard to identify differences between *RAS* alleles that drive tumorigenesis. For example, exogenous gene expression is a commonly used tool, yet *RAS* gene dosage has been shown to effect signaling (Zhang et al. 2007), localization (Nan et al. 2015) and *in vivo* functionality (Omerovic et al. 2007; Xu et al. 2013). The biological consequences of mutant RAS expression also differ based on the isoform (H-, K- or N-RAS) and cell-type examined (Yan et al. 1998; Voice et al. 1999; Haigis et al. 2008; Burd et al. 2014; Kong et al. 2016). Therefore, it is essential to assess the differences between endogenous RAS mutants under physiologically relevant conditions.

Here, we report the development of a suite of eight NRAS*-*mutant mouse alleles, each of which enables the conditional expression of a distinct NRAS mutant from the endogenous gene locus. Crossing these alleles to a melanocyte-specific Cre, we find that the melanomagenic potential of NRAS mutants parallels the frequency of these alleles in human melanoma. Further, we link the melanomagenic potential of NRAS mutants to enhanced RAF dimerization and heightened MEK/ERK MAP kinase pathway signaling.

## RESULTS

### The tumorigenic potential of NRAS mutants parallels allelic frequency in human melanoma

We used CRISPR-Cas9 to zygotically modify the *Nras* mutation in Tyr::CreER^T2^; *LSL-Nras*^*61R*/*R*^ (*TN*^*61R*/*R*^) mice (**Fig. S1A-B, S2A**; (Burd et al. 2014; Hennessey et al. 2017)). This process yielded eight new mouse models in which Cre recombinase triggers the melanocyte-specific expression of a modified *Nras* gene from the endogenous locus: *TN*^*61K*/*K*^, *TN*^*61L*/*L*^, *TN*^*61H*/*H*^, *TN*^*61P*/*P*^, *TN*^*61Q*/*Q*^, *TN*^*12D*/*D*^, *TN*^*13D*/*D*^ and *TN*^*13R*/*R*^. Each *LSL-Nras* allele was sequenced and functionally validated in murine embryonic fibroblasts (MEFs) (**Fig. S1C-E, S2B-D**). Founder animals were then backcrossed two generations to *TN*^*61R*/*R*^ mice to limit any off-target effects of CRISPR-Cas9.

We employed this suite of *TN* mice to determine if *NRAS* oncogenes common to human melanoma (**Fig. 1A**) could drive melanocyte transformation better than those present in other tumor types. Experimental *TN*^*61X*/*X*^ cohorts were generated by intercrossing Tyr::CreER^T2^ transgenic mice carrying one *LSL-Nras*^*61R*^ and one *LSL-Nras*^*61X*^ allele, where X = K, L, H, P or Q (**Fig. S1F**). The resulting offspring were topically treated with 4-hydroxytamoxifen (4-OHT) on post-natal days 1 and 2 to drive CreER^T2^-mediated excision of the *LSL* transcriptional stop sequence and initiate expression of each *Nras* variant (**Fig. S1G**). The mice were then subjected to a single, 4.5 kJ/m^2^ dose of ultraviolet B (UVB) irradiation on post-natal day 3 to mimic the role of sunlight in melanoma formation (**Fig. S1G**) (Hennessey et al. 2017).

**Figure 1:**
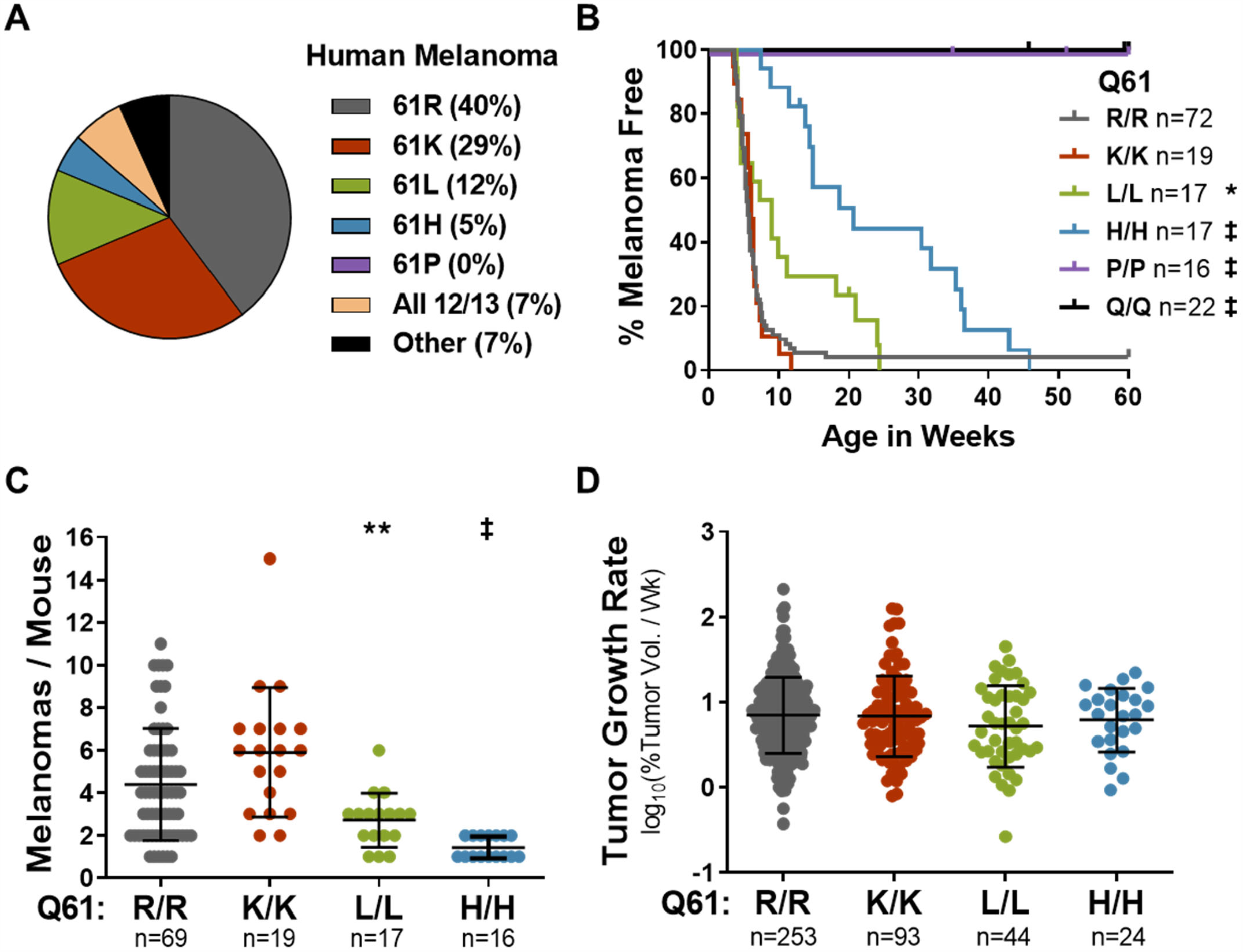
Frequency of NRAS mutants in human melanoma parallels tumorigenic potential in mice. **A**, Frequency of *NRAS* mutations in the TCGA PanCancer Atlas dataset for human cutaneous melanoma. **B-D**, Melanoma-free survival (B), total tumor burden (C) and tumor growth rates (D) for mice expressing the indicated melanocyte-specific NRAS mutants. Log-rank (Mantel-Cox) (B) or ANOVA (C-D) with a Dunnet T3 multiple comparisons test was used to compare measurements between each genotype and *TN*^*61R*/*R*^. Adjusted p-values for all comparisons can be found in **Table S1A. \*** p< 0.05, **\*\*** p< 0.01, ‡ p< 0.0001.

Spontaneous melanomas formed more rapidly and frequently in *TN*^*61R*/*R*^ and *TN*^*61K*/*K*^ mice than in *TN*^*61L*/*L*^ or *TN*^*61H*/*H*^ animals, and no tumors were detected in the *TN*^*61P*/*P*^ and *TN*^*61Q*/*Q*^ models (**Fig. 1B-C, Table S1A**). These differences were not due to litter-specific effects as the onset, burden and growth rates of *TN*^*61R*/*R*^ tumors did not differ between experimental cohorts or male and female mice (**Fig. S1H-L**). Melanoma growth rates were similar regardless of genotype (**Fig. 1D, Table S1A**), leading to overall survival rates which paralleled the tumor onset for each *TN*^*61X*/*X*^ model (**Fig. S1M**). Hematoxylin and eosin stained tumor sections from each NRAS model exhibited similar morphology and invasiveness (data not shown). UVB light cooperated equally with each NRAS mutant to enhance tumor onset and burden, revealing that differences in the melanoma-driving capabilities of each variant are independent of UVB carcinogenesis (**Fig. S1N-Q, Table S1B**).

Our results in the *TN*^*61X*/*X*^ models and the rarity of codon 12/13 mutants in human melanoma suggested that *TN*^*12D*/*D*^, *TN*^*13D*/*D*^ and *TN*^*13R*/*R*^ mice would not develop tumors. To test this hypothesis, we generated experimental colonies by breeding mice homozygous for each codon 12 or 13 allele in our series. *TN*^*12D*/*D*^ and *TN*^*13D*/*D*^ mice did not succumb to melanoma after 60 weeks of observation (**Fig. S2E-F**). By contrast, *TN*^*13R*/*R*^ mice did form melanomas, albeit with lower efficiency than even the weakest melanoma-forming codon 61 model, *TN*^*61H*/*H*^. These data, summarized in **Table S1C**, establish differences in the ability of oncogenic NRAS mutants to initiate melanoma formation and provide a plausible explanation for the prevalence of *NRAS*^*61R*^ and *NRAS*^*61K*^ mutations in human melanoma.

### NRAS proteins with compromised GTPase activity facilitate NRAS^61R^-dependent melanomagenesis

In RAS-driven malignancies, the complementary wild-type allele is thought to suppress tumorigenesis driven by the mutationally-active oncoprotein (Bremner and Balmain 1990; Zhang et al. 2001; To et al. 2013; Kong et al. 2016). However, studies examining the interaction between alleles of differing oncogenic potential are lacking and could shed light on the functional interplay between RAS molecules. To explore this question, we compared the tumor onset, burden and overall survival of homozygous (*TN*^*61R*/*R*^, *TN*^*61X*/*X*^) and heterozygous (*TN*^*61X*/*R*^) mice from each of our experimental cohorts. The melanoma phenotypes of heterozygous mice were generally intermediate to those observed in homozygous *TN*^*61R*/*R*^ and *TN*^*61X*/*X*^ animals from the same cohort (**Fig. 2A-D, Table S1C**). However, NRAS^61R^ was unable to drive melanoma formation in the presence of wild-type NRAS^61Q^ (**Fig. 2E**). These data reveal that the additive effect of *Nras* alleles in melanoma is dependent upon intact GTPase activity.

**Figure 2:**
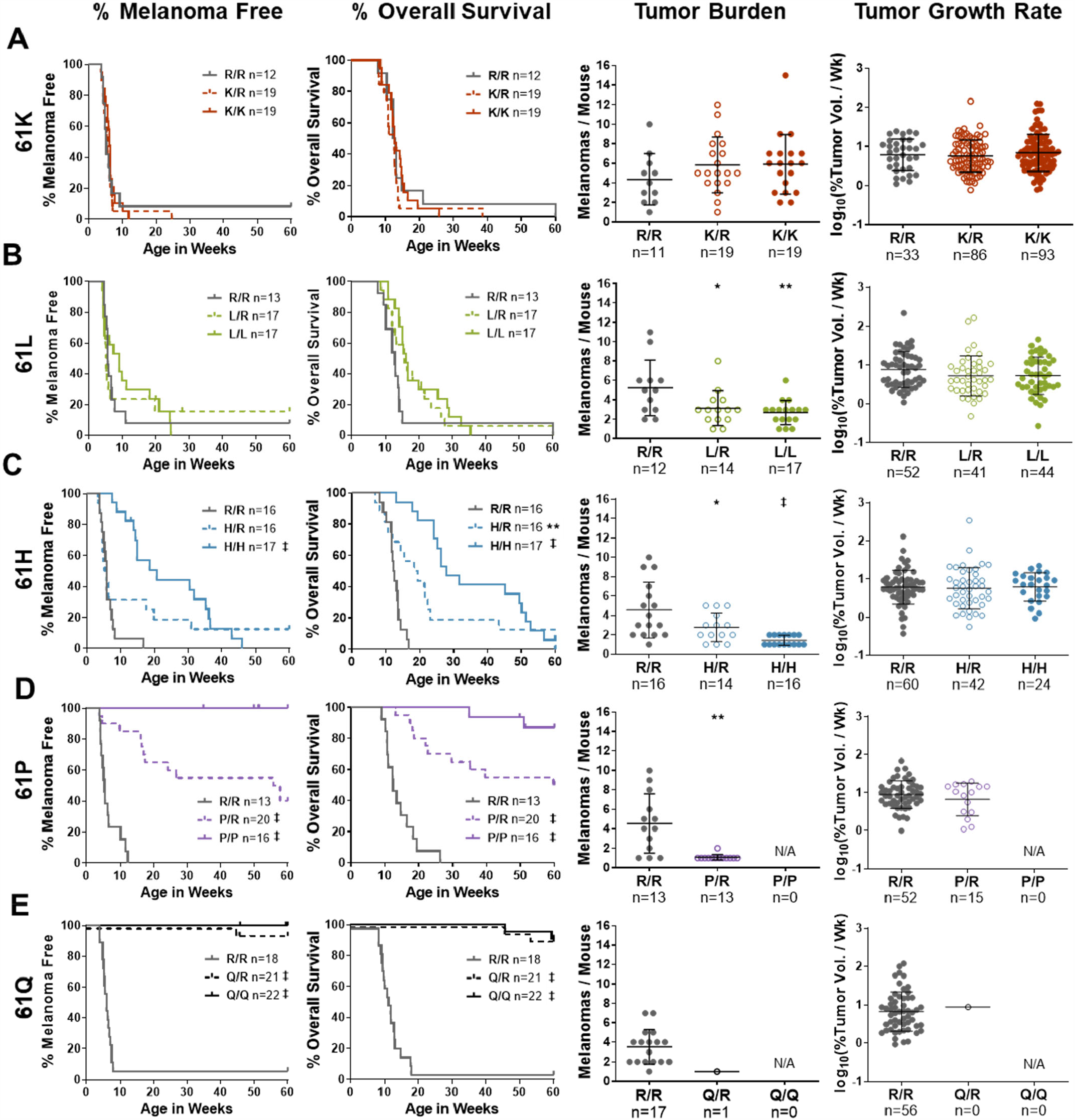
Combining codon 61 alleles defective in GTPase activity results in an intermediate melanoma phenotype. **A-E**, Melanoma-free survival, overall survival, tumor burden and tumor growth rates for the following treatment cohorts: (A) *TN*^*61K*/*R*^, (B) *TN*^*61L*/*R*^, (C) *TN*^*61H*/*R*^, (D) *TN*^*61P*/*R*^, and (E) *TN*^*61Q*/*R*^. In A-E, the phenotype of *TN*^*61R*/*R*^ mice was compared to *TN*^*61X*/*X*^ and *TN*^*61X*/*R*^ animals. Log-rank (Mantel-Cox) tests were used to compare survival. ANOVA with a Dunnet T3 multiple comparisons test was used to compare tumor burden and growth. **\*** p< 0.05, **\*\*** p< 0.01, ‡ p< 0.0001.

### Tumorigenic NRAS proteins drive increased proliferative signaling

We performed RNA sequencing on MEFs derived from our melanomagenic (61R/R and 61H/H) and non-melanomagenic (61P/P and 61Q/Q) *TN* models to identify mechanistic differences between NRAS mutants. RNA was isolated from passage five MEFs six days after NRAS induction. We first compared the transcriptome of cells expressing each NRAS mutant (61P, 61H or 61R) to those expressing the wild-type protein (61Q). Fewer transcripts were differentially expressed in NRAS^61P/P^ than NRAS^61R/R^ or NRAS^61H/H^ MEFs as compared to wild-type controls (NRAS^61Q/Q^) (2413, 4137, and 5221 transcripts, respectively; **Fig S3A, C-E; Table S2A-C**). Principal component analysis (PCA) showed that MEFs expressing melanomagenic NRAS mutants (NRAS^61R/R^ or NRAS^61H/H^) clustered separately from those expressing NRAS^61P/P^ or NRAS^61Q/Q^ (**Fig S3B**). MEFs expressing NRAS^61P/P^, NRAS^61R/R^ or NRAS^61H/H^ were enriched for transcripts associated with E2F, MYC and the G2M checkpoint as compared to NRAS^61Q/Q^ MEFs; however, only NRAS^61R/R^ MEFs showed an enrichment for genes associated with heightened RAS signaling (**Fig S3F-H)**.

RAS mutants exhibit distinct biochemical properties which, based on our *in vivo* data, may be essential for melanomagenesis. Therefore, we next sought to identify transcriptional differences between MEFs expressing melanomagenic and non-melanomagenic NRAS mutants. Over 1,600 transcripts differed between MEFs expressing the non-melanomagenic NRAS^61P/P^ and melanomagenic NRAS^61R/R^ and NRAS^61H/H^ mutants (1631 and 1844, respectively; **Fig. 3A-B; Table S3A-C**). MEFs expressing melanomagenic NRAS mutants were enriched for transcripts associated with MYC and KRAS signaling, but also exhibited upregulation of inhibitors of the MAPK pathway, such as DUSP6 and SPRY2 (**Fig. 3C-D, Fig S3I-J**). Despite upregulation of these inhibitors, MEFs and cutaneous melanocytes expressing melanomagenic NRAS mutants showed heightened proliferation as determined by EdU incorporation (**Fig. 3E, Fig. S3K-L, Table S4A**). These observations highlight differences in the transcriptomes elicited by NRAS mutants that are, and are not, capable of initiating melanoma. Specifically, melanoma-driving NRAS mutants are able to subvert cellular feedback mechanisms that limit MAPK activity. This is consistent with data from human melanomas where *DUSP6* and *SPRY2* are upregulated as compared to normal skin (**Fig. S3M-N**).

**Figure 3:**
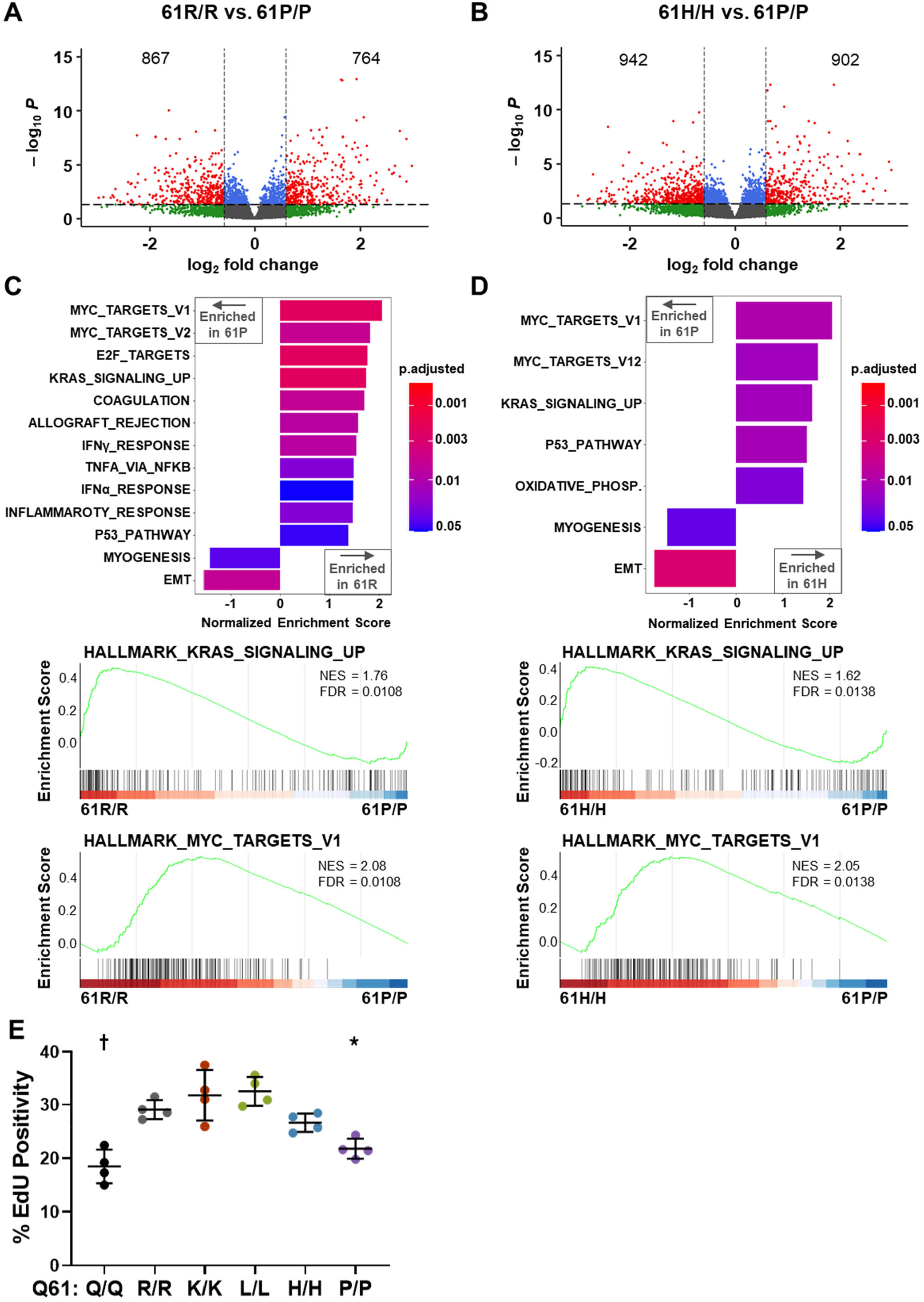
Differential regulation of the RAS-Myc axis by tumorigenic and non-tumorigenic NRAS mutants. **A-B**, Volcano plot depicting differentially expressed genes in MEFs expressing NRAS^61R/R^ versus NRAS^61P/P^ (A) or NRAS^61H/H^ versus NRAS^61P/P^ (B). **C-D**, (top) Bar plot showing the differential enrichment of Hallmark gene sets (p-adjusted < 0.05) in MEFs expressing NRAS^61R/R^ versus NRAS^61P/P^ (C) or NRAS^61H/H^ versus NRAS^61P/P^(D). (bottom) Enrichment plots generated from differentially expressed genes associated with the Hallmark gene sets: KRAS_SIGNALING_UP or MYC_TARGETS_V1. **E**, Flow cytometric analysis of EdU labeling in NRAS-mutant MEFs. Each dot represents one biological replicate. ANOVA with a Tukey post-test was used to compare data from *Nras*^*61R*/*R*^and *Nras*^*61X*/*X*^ cells. Adjusted p-values for all comparisons can be found in **Table S4A. \*** p< 0.05, † p< 0.001.

To test the idea that MAPK signaling is sustained in the presence of melanomagenic NRAS mutants, we analyzed ERK and AKT activation in MEFs and tumors from the *TN*^*61X*/*X*^ models. NRAS was induced with adenoviral Cre in MEFs and six days later, the cells were placed in serum-free media for 4 hours prior to protein isolation. MEFs expressing NRAS^61R/R^ or NRAS^61K/K^ had higher levels of phospho-ERK than MEFs expressing NRAS^61L/L^, NRAS^61H/H^, NRAS^61P/P^ or NRAS^61Q/Q^ (**Fig. 4A, Table S4A**). However, signals downstream of RAS were not similarly amplified by the oncogenic alleles, as activation of the PI3K/AKT signaling pathway was similar regardless of *Nras* genotype (**Fig. 4A**). The same mutation-specific effects on NRAS signaling were observed in melanomas from our *TN*^*61R*/*R*^ and *TN*^*61K*/*K*^ mice as compared to tumors derived from the *TN*^*61L*/*L*^ and *TN*^*61H*/*H*^ models (**Fig. 4B, Table S4A**). These results suggest that mutation-specific differences in MAPK activation dictate the melanomagenic potential of NRAS in mice.

**Figure 4:**
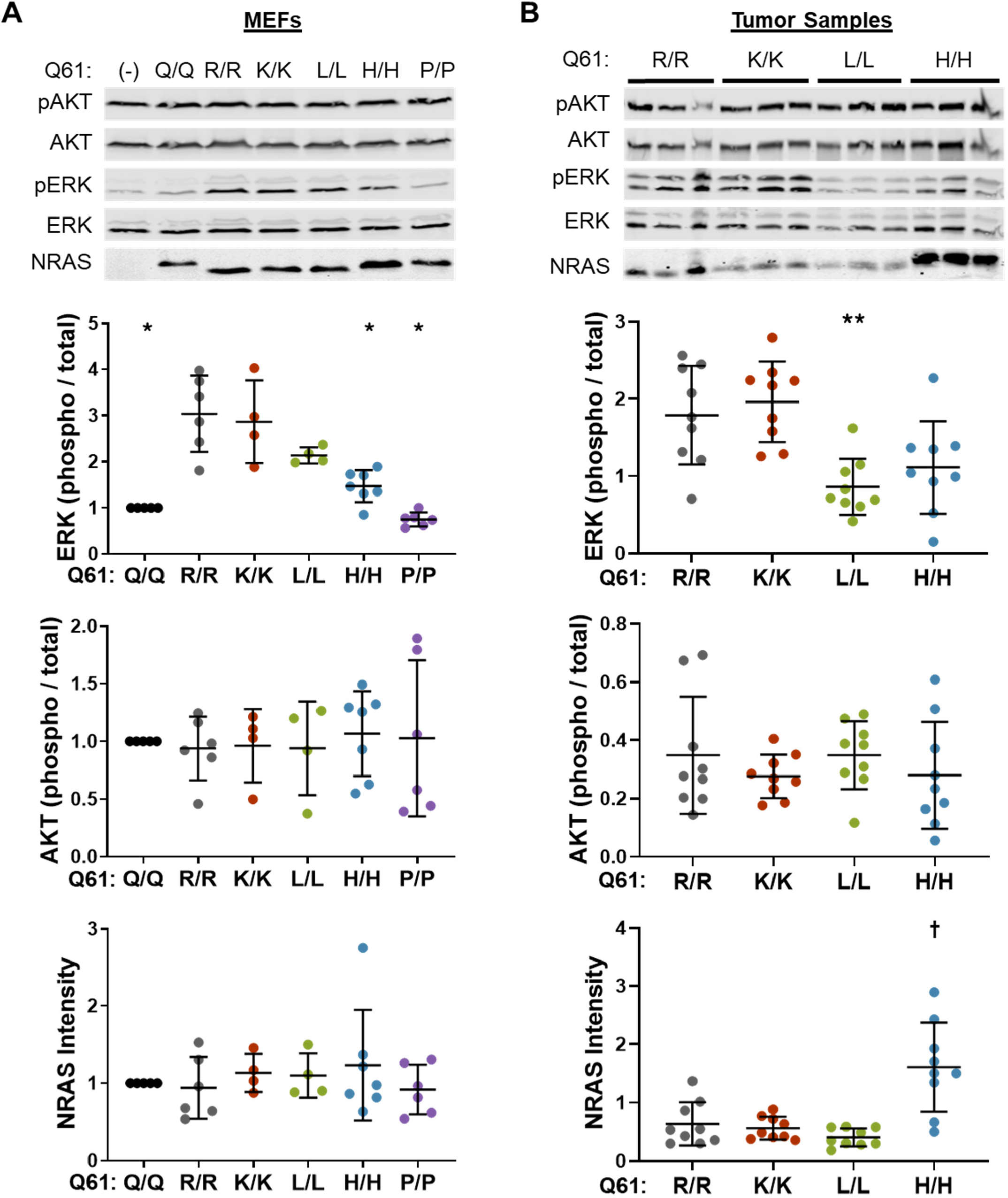
MAPK pathway activation parallels the tumorigenic potential of oncogenic NRAS mutant. **A-B**, Immunoblot of protein lysates isolated from MEFs (A) or murine melanomas (B) expressing the indicated NRAS mutants. Dot plots showing the quantification of ERK activation, AKT activation or NRAS expression. Each dot represents one biological replicate. ANOVA with a Dunnet T3 multiple comparisons test was used to compare data from each codon 61 mutant to MEFs expressing NRAS^61R/R^ and an ANOVA with a Tukey’s multiple comparison test was used to compare data from each codon 61 mutant to protein lysate isolated from mice expressing NRAS^61R/R^. Adjusted p-values for all comparisons can be found in **Table S4A. \*** p< 0.05, **\*\*** p< 0.01, † p< 0.001.

### Melanomagenic NRAS mutants promote RAF dimerization to drive enhanced MAPK signaling

Mutationally-active RAS proteins stimulate signaling through the RAF>MEK1/2>ERK1/2 pathway through both direct and indirect mechanisms (Zhou et al. 2016). Mutant RAS can indirectly activate MAPK through the allosteric regulation of SOS1, which in turn promotes GTP loading on wild-type RAS isoforms (Jeng et al. 2012). To determine if melanomagenic NRAS mutants promote higher levels of MAPK signaling via this indirect mechanism, we used lentiviral shRNAs to knockdown *Sos1* or *Hras* and *Kras* in NRAS^61R/R^, NRAS^61P/P^ and NRAS^61Q/Q^ MEFs. These knockdowns had no effect on MAPK activation regardless of the *Nras* allele present, ruling out the possibility that melanomagenic alleles drive heightened MAPK signaling through indirect activation of wild-type RAS (**Fig. 5A, Table S4B**). Knockdown of *Nras* served as a positive control and reduced MAPK pathway activation in MEFs expressing NRAS^61R/R^ (**Fig. 5A**).

**Figure 5:**
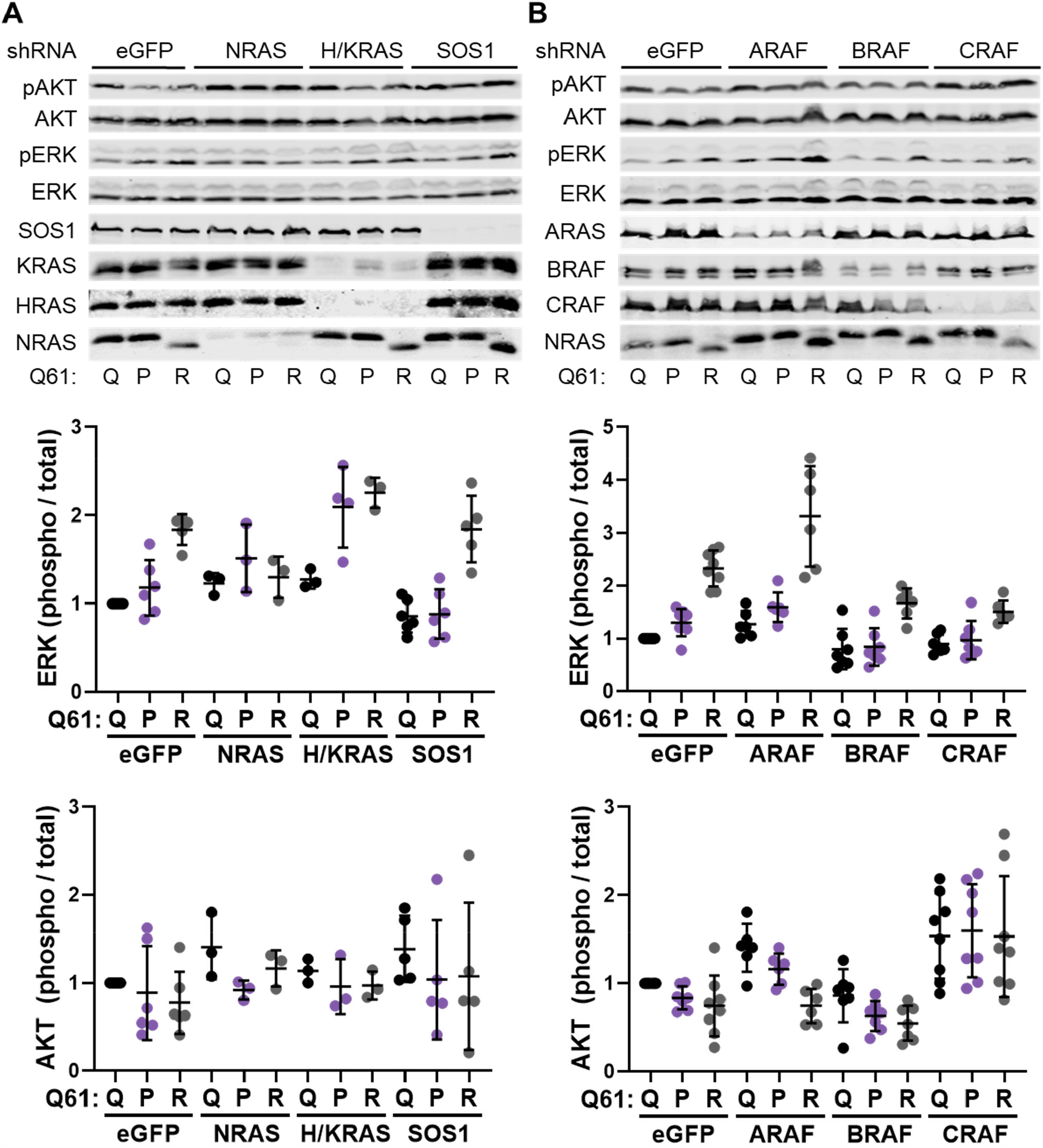
Oncogenic NRAS mutants mediate differential MAPK activation via a RAF-dependent mechanism. **A-B**, Representative immunoblots of AKT and ERK activation in homozygous MEF cell lines treated with shRNAs targeting *Nras, Hras* and *Kras*, or *Sos1* (A) or *Araf, Braf* or *Craf* (B). Dot plots show the quantification of AKT and ERK activation in biological replicates, each represented by a single dot. Adjusted p-values were generated using ANOVA with a Dunnet T3 multiple comparisons test. Adjusted p-values can be found in **Table S4B-C**.

RAS isoforms and KRAS mutants have been shown to differentially engage RAF monomers in exogenous expression systems (Terrell et al. 2019). Thus, we postulated that melanomagenic NRAS mutants might activate RAF better than non-melanomagenic mutants in our endogenous expression system. Knockdown of *Braf* or *Craf* using lentiviral shRNA partially reduced MAPK activation in NRAS^61R/R^, NRAS^61P/P^ and NRAS^61Q/Q^ MEFs (**Fig 5B, Table S4C**). However, the most dramatic effect was observed with *Craf* loss in NRAS^61R/R^ MEFs (p < 0.01 vs NRAS^61R/R^ eGFP) (**Fig. 5B**). *Araf* knockdown, by contrast, enhanced ERK activation in the NRAS^61R/R^ MEF line (**Fig. 5B**). To further confirm these results, we developed an adenoviral NanoBiT system to measure the homo- and hetero-dimerization of ARAF, BRAF and CRAF in live cells (**Fig. 6A**). We induced NRAS expression in MEFs from each of our *LSL-Nras*^*61X*/*X*^ alleles and then infected the cells with adenovirus encoding BRAF-LgBiT and BRAF-SmBiT (**Fig. 6B**), CRAF-LgBiT and CRAF-SmBiT (**Fig. 6C**) or BRAF-LgBiT and CRAF-SmBiT (**Fig. 6D**). These results showed that the ability of NRAS mutants to drive melanoma *in vivo* parallels the induction of BRAF-BRAF and BRAF-CRAF dimers *in vitro* (**Table S4D**). Together, our findings suggest a model in which NRAS mutants capable of initiating melanoma formation promote enhanced BRAF dimerization, leading to heightened MAPK signal transduction (**Fig. 7**). Notably, the lack of PI3K activation across the alleles (**Fig. 4**) suggests that melanoma-initiating NRAS mutants exhibit specific effector preferences.

**Figure 6:**
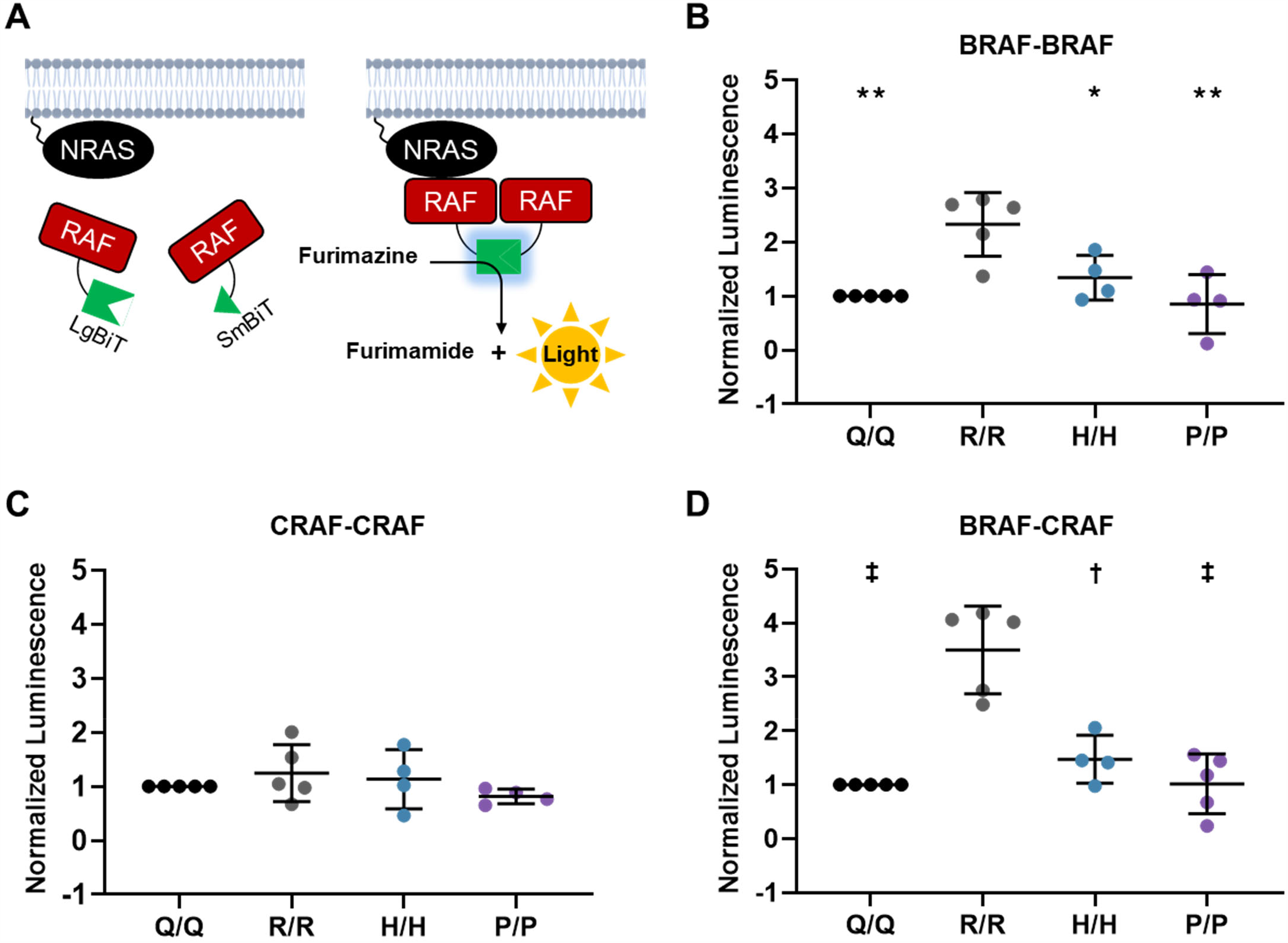
Melanomagenic NRAS mutants drive enhanced RAF dimerization. **A**, Schematic representation of the RAF NanoBiT assay in which each RAF isoform is tagged with either LgBiT or SmBiT. **B-D**, Dot plot of normalized luminescence intensity in *TN*^*61X*/*X*^ MEFs infected with adenovirus expressing BRAF-LgBiT and BRAF-SmBiT (B), CRAF-LgBiT and CRAF-SmBiT (C), or BRAF-LgBiT and Craf-SmBiT (D). Luminescence intensity was normalized to crystal violet staining for each well. ANOVA with a Dunnet T3 multiple comparisons test was used to compare luminescence intensity from each codon 61 mutant to MEFs expressing NRAS^61R/R^. Adjusted p-values for all comparisons can be found in **Table S4D. \*** p< 0.05, **\*\*** p< 0.01, † p< 0.001, ‡ p< 0.0001.

**Figure 7:**
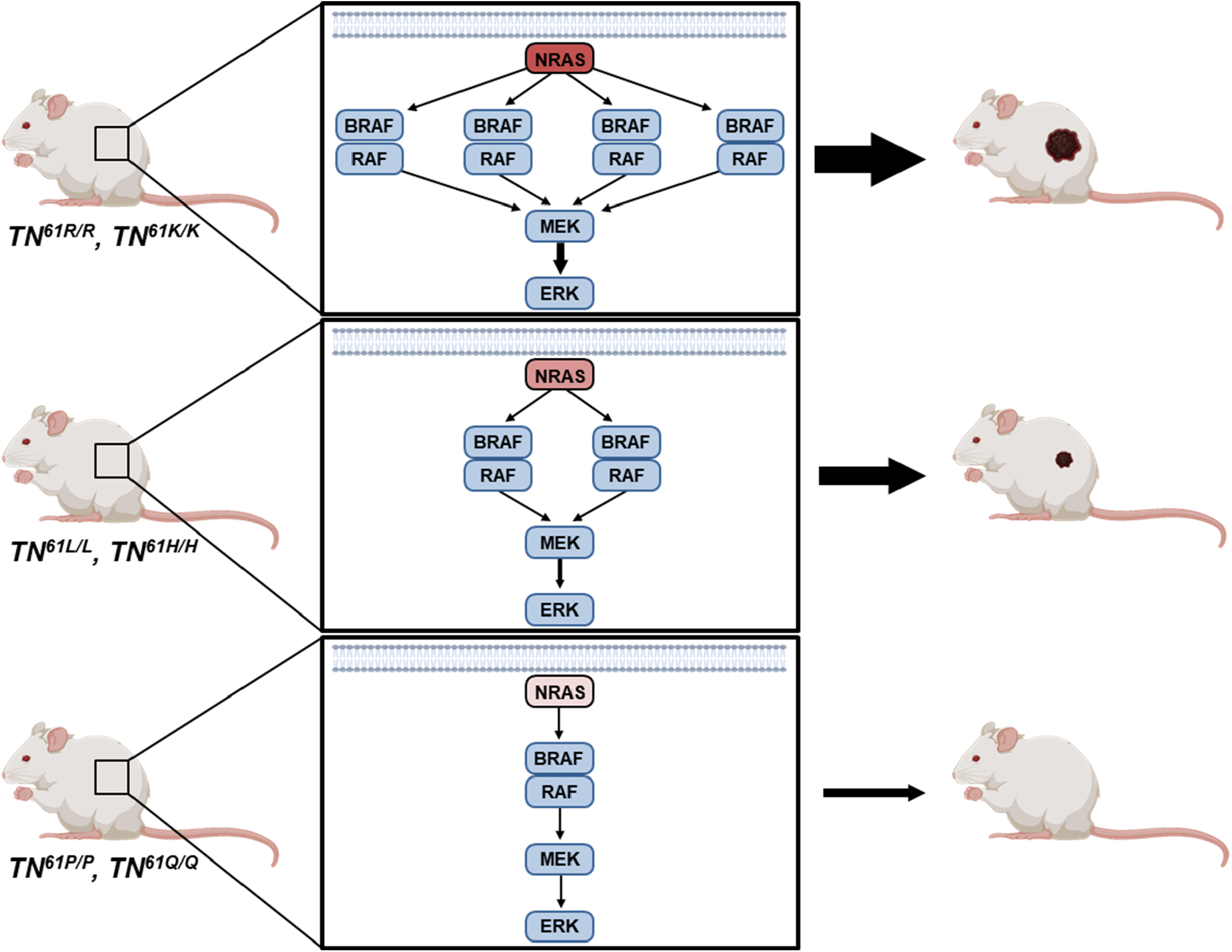
Differential RAF engagement explains variances in the ability of oncogenic NRAS mutants to initiate melanoma formation.

## DISCUSSION

Our data establish that functional differences, rather than mutagenic preferences, explain the frequency of NRAS mutants in human melanoma. This observation opens the door for new therapeutic and preventative strategies targeting functions exclusive to melanomagenic NRAS mutants. Putting forth this versatile suite of inducible, endogenous *Nras* alleles will support the broader scientific community with similar endeavors in a variety of cancer types and permit the investigation of mutant-specific drug sensitivities.

Previous publications highlight differences in the tumorigenic potential of *RAS* codon 12 and 61 mutations in pancreatic cancer, lung cancer, leukemia and melanoma (Burd et al. 2014; Park et al. 2015; Kong et al. 2016; Zhou et al. 2020). However, these results might be predicted as codon 12 and 61 mutants distinctly alter GTP activity. Codon 12 and 13 mutants prevent GAP-mediated catalysis of GTP hydrolysis (Adari et al. 1988), whereas codon 61 alterations abolish intrinsic RAS GTPase activity (Frech et al. 1994). On the other hand, minimal differences are observed in the GTP hydrolysis rates between RAS codon 61 mutants (Novelli et al. 2018),raising the question of why certain codon 61 mutants would be more prevalent in melanoma. We explored this question using a new suite of melanoma GEMMs that enable the conditional and melanocyte-specific expression of eight different NRAS mutants from the endogenous gene locus. Using these mice, we show that NRAS codon 61 variants possess distinct melanomagenic potential when expressed at endogenous levels *in vivo* (**Fig. 1B**). Therefore, it is unlikely that selective carcinogenesis determines prevalence of specific NRAS mutants in human melanoma.

In human cancers, *RAS* mutations typically exist in a heterozygous state. Here, the effect of the cognate wild-type allele is often tumor suppressive (Zhou et al. 2016). Therefore, loss or downregulation of the wild-type allele is seen in many RAS-driven tumors (Bremner and Balmain 1990; Zhang et al. 2001; To et al. 2013). Still, isoform- and tissue-specific functions prevent these results from being extrapolated across all RAS-driven cancers. In our models, NRAS^61R^ was unable to drive tumorigenesis in the presence of a wild-type NRAS^61Q^ (**Fig. 2E**), whereas an additive tumorigenic potential was observed when NRAS^61R^ was paired with other, constitutively active NRAS variants (**Fig. 2A-D**). While these data suggest a tumor suppressive role of wild-type NRAS in melanoma, we are unable to rule out the possibility that gene dosage is an important determinant of tumorigenic potential as previously reported in murine models of acute myeloid leukemia (Xu et al. 2013). Future studies to address this question will require that melanoma formation is assessed in a *TN*^*61R*/*R*^ model that has been crossed to a conditional *Nras* knockout allele.

Here we show that the ability to enhance MAPK signaling is a key feature of melanomagenic NRAS mutants. MAPK signaling plays a pivotal role in the evolution of human melanoma, with elevated activity occurring early and strengthening throughout disease progression (Shain et al. 2018). Notably, non-melanomagenic mutants, such as NRAS^12D^, are commonly detected in combination with *NRAS* amplification or activating mutations in other components of the MAPK pathway (Cerami et al. 2012; Gao et al. 2013). Studies in a mouse model expressing NRAS^12D^ support this notion, showing that the mutant drives cutaneous melanoma only in the presence of a kinase-dead BRAF capable of inducing paradoxical RAF activation (Heidorn et al. 2010). These observations support a critical role for MAPK signaling in melanoma initiation (Shain et al. 2018). We observe differences in MAPK activation shortly after the induction of NRAS expression in our models and show that these differences persist throughout tumorigenesis (**Fig. 4**). These data suggest that the melanomagenic potential of a given NRAS mutant is determined by its ability to sustain MAPK activation. This observation may also explain why melanoma progression is more rapid in UV-irradiated models (**Fig. S1N-Q, Table S1B**) (Viros et al. 2014; Hennessey et al. 2017; Pérez-Guijarro et al. 2017) as UV light stimulates surrounding keratinocytes to secrete paracrine growth factors and cytokines that may further augment MAPK signaling (Wang et al. 2016). Nevertheless, the fact that negative regulators of the MAPK pathway are elevated both in our mouse models and in human melanomas makes it clear that MAPK signaling must be carefully balanced during disease onset. Perturbing this balance, in one direction or the other, could be key to melanoma prevention.

Our data support a mutant- and disease-specific approach to targeting RAS-driven cancers. Complete blockade of NRAS^61R^ or NRAS^61K^ may not be necessary if the functional properties of these alleles could be shifted toward a more NRAS^12D^- or NRAS^61P^-like phenotype or conformation. Our data suggest that drugs shifting effector engagement would be effective in preventing NRAS-mutant melanoma. It is likely that this approach will also be broadly applicable. Isoform and mutation-specific differences in RAS-RAF interactions have been reported in resonance energy transfer and Co-IP experiments using exogenously expressed, tagged constructs (Terrell et al. 2019). Building upon these observations, our findings establish differences in the ability of endogenous NRAS mutants to drive the formation of specific RAF dimers in live cells (**Fig. 6**). As such, the selective disruption of RAF dimers deemed essential for disease progression could lead to viable prevention and treatment strategies with reduced toxicity.

## METHODS AND MATERIALS

### Murine alleles and husbandry

Animal work was performed in compliance with protocols approved by The Ohio State Institutional Care and Use Committee (Protocol #2012A00000134). The *LSL-Nras*^*61R*^ allele and *TN* model were previously described and have been backcrossed >7 generations to C57BL/6J (Burd et al. 2014) (MMRRC #043604-UNC). Other *LSL-Nras*^*61X*^ alleles were created via zygotic gene editing with CRISPR-Cas9 technology (gRNA and homology oligo sequences provided in **Table S5A**). Codon 61 alleles were generated from *TN*^*61R*/*R*^ homozygous zygotes, whereas codon 12 and codon 13 alleles were generated from *TN*^*61Q*/*Q*^ homozygous zygotes. Targeting was verified in the resulting offspring by Sanger Sequencing (primers provided in **Table S5A**). During this process, a silent G/A mutation was discovered in the 3^rd^ nucleotide of codon 15 of the *LSL-Nras*^*12D*^ and *LSL*-*Nras*^*13R*^ alleles. Each allele was backcrossed two generations to *TN*^*61R*/*R*^ mice prior to beginning experiments.

### In vivo Cre induction and UV exposure

NRAS expression was initiated as described (Burd et al. 2014) by applying 20 mM 4-hydroxytamoxifen (4-OHT) to the backs of neonatal pups on post-natal days one and two. On post-natal day three, animals were subjected to a single, 4.5 kJ/m^2^ dose of ultraviolet B (UVB) using a fixed position 16W, 312 nm UVB light source (Spectronics #EB-280C). [See (Hennessey et al. 2017) for additional information.]

### Outcome monitoring and histopathology

Mice from each cohort were randomly numbered and blindly monitored three times a week for tumor formation. Upon detection, melanomas were measured three times per week and tumor size (width x length (mm)) was recorded using calipers. Tumors reaching protocol exclusion criteria were harvested. A portion of each primary tumor was fixed in 10% neutral buffered formalin and the rest was flash-frozen for protein extraction. Formalin-fixed samples were paraffin embedded, sectioned (4 µm) and stained with hematoxylin and eosin (H&E). Stained tumor sections were evaluated using an Olympus BX45 microscope with attached DP25 digital camera (B&B Microscopes Limited, Pittsburgh, PA) by a veterinary pathologist certified by the American College of Veterinary Pathologists (K.M.D.L.).

### Isolation and culture of primary mouse embryonic fibroblasts

Mouse embryonic fibroblasts (MEFs) were generated from E13.5 embryos using manual homogenization and trypsinization. Dissociated cells were cultured in Dulbecco’s modified eagle medium (DMEM), supplemented with 10% fetal bovine serum (FBS), 1% penicillin-streptomycin and 1% glutamine. MEF lines were passaged when confluency reached 70-80% in a 10 cm tissue culture dish.

### In vitro induction of NRAS expression

MEFs were seeded at equal density in 10 cm tissue culture plates. The following day, these cultures were washed with PBS and placed in DMEM containing 0.5% FBS, 1% penicillin-streptomycin and 1% glutamine. Adenovirus expressing Cre recombinase conjugated to eGFP or eGFP alone (Ad5-CMV-Cre-eGFP, Ad5-CMV-eGFP; Baylor College of Medicine Vector Development Laboratory, Houston, TX) was added to the cultures for 16 hours at an MOI of 4000:1 (viral particles : cells). After infection, cells were allowed to recover for at least 72 hours in DMEM containing 10% FBS, 1% penicillin-streptomycin and 1% glutamine prior to analysis. Allelic recombination was confirmed through genomic PCR as described (Burd et al. 2014).

### Immunoblotting

Frozen tumors (10-15 mg) were homogenized using a liquid nitrogen-cooled mortar and pestle. Homogenized tumor tissue and pelleted cell lines were lysed in RIPA (25mM Tris pH 7.4, 150mM NaCl, 1% IGEPAL, 0.1% SLS) supplemented with protease inhibitor cocktail (Sigma P8340), calyculin A (CST 9002S) and Halt phosphatase inhibitor cocktail (Thermo Fisher 78420). Equal protein concentrations, as determined by Bradford Assay (Bio-Rad #5000006), were run on an SDS-PAGE gel and transferred to PVDF (Sigma IPFL00010). PVDF membranes were blocked in 5% milk-PBS and then probed with one of the following primary antibodies: ERK1/2 (1:1000, CST 4696S), phospho-ERK1/2 (1:1000, CST 9101S), AKT (1:1000, CST 2920), phospho-AKT (1:1000, CST 9271), NRAS (1:250, Abcam ab77392), HRAS (1:1000, Abcam ab32417), KRAS (1:1000, Sigma WH0003845M1), SOS1 (1:1000, CST 5890), ARAF (1:1000, CST 4432P), BRAF (1:500, Santa Cruz sc-5284), or CRAF (1:500, CST 12552). Secondary antibodies were diluted in 5% BSA 1x PBST as follows: anti-goat (1:15000. LI-COR 926-32214), anti-mouse (1:15000, LI-COR 926-68070, LI-COR 926-32210), or anti-rabbit (1:15000, LI-COR 926-68071, LI-COR 926-32211). Membranes were imaged on a LI-COR Odyssey CLx system and quantified using Image Studio software (LI-COR Biosciences).

### RNA-Sequencing

NRAS expression was induced in passage 3 MEFs using Ad5-CMV-Cre-eGFP as described for our *in vitro* studies. The cells were then cultured for 6 days prior to RNA isolation using the ZR-Duet DNA/RNA MiniPrep Plus Kit (Zymo D7003). RNA quality and concentration were confirmed on an Agilent TapeStation and Life Technologies Qubit. RNA was prepared for sequencing through ribosomal depletion using Illumina Ribo-Zero chemistry followed by library preparation using Illumina TruSeq Total RNA Stranded Library Prep Kit. RNA was sequenced on an Illumina HiSeq4000 with 150 base-pair, paired-end reads. Raw data files are deposited in the NCBI Gene Expression Omnibus (GEO) under accession #GSE162124.

RNA reads were aligned to build 38 of the mouse genome (mm10) using STAR (Dobin et al. 2013), duplicates marked using PICARD (version 2.17.11) (http://broadinstitute.github.io/picard/) and a gene count matrix generated by featureCounts (Liao et al. 2014). Differential gene expression analysis was performed using DESeq2 (*p*-adjusted < 0.05) (Love et al. 2014). Gene set enrichment analysis used the DOSE algorithm within the GSEA function of the clusterProfiler package (Yu et al. 2012; Yu et al. 2015) to probe gene sets from the molecular signatures database Hallmark collection (Liberzon et al. 2015).

### Flow cytometric analysis of EdU labeling

Passage three *TN*^*61X*/*X*^ MEFs were infected with Ad5-CMV-Cre-eGFP to induce NRAS expression as described for our *in vitro* studies and then cultured for five days. MEFs were then incubated in DMEM containing 1% penicillin-streptomycin and 1% glutamine for five hours prior to adding 0.01 mM 5-ethynyl-2-deoxyuridine (EdU) to the media. MEFs were labeled with EdU for an additional five hours and then harvested and fixed with 4% paraformaldehyde. Fixed cells were permeabilized with saponin in 1% BSA 1x PBS. Click-iT chemistry was used to label the incorporated EdU with Chromeo 642. Here, the cells were incubated for 30 minutes in Click-iT reaction cocktail containing 2 mM CuSO_4_, 50 mM ascorbic acid and 50 nM Chromeo 642 azide dye (Active Motif 15288) diluted in 1x PBS. 10,000 cells per sample were analyzed on a BD LSR Fortessa flow cytometer and the percentage of EdU positive cells was determined using FlowJo software. Specifically, the initial population of MEFs was selected by gating based on FSC-A by SSC-A (**Fig S4**, top). Next, cell doublets were removed by gating for single cells in a FSC-H by FSC-A plot (**Fig S4**, middle). Finally, a histogram of counts by APC-A intensity was used to determine the percent of EdU positive cells in each population of MEFs (**Fig S4**, bottom).

### EdU labeling of melanocytes in vivo

Neonatal pups were induced to express NRAS and treated with UVB irradiation as described above. EdU (0.041 mg/kg) was administered to mice on post-natal day 10 via intraperitoneal injection. Two hours later, mice were euthanized and the dorsal skin collected. Samples were fixed in 10% neutral buffered formalin for 24 hours, embedded in paraffin and cut into 5 µm sections. Slides of each section were deparaffinized and rehydrated and then Click-iT chemistry was used to label the incorporated EdU with Chromeo 642 as described above. Antigen retrieval was performed using Dako Antigen Retrieval Solution (Agilent S169984-2) followed by blocking with Dako Protein Block (Agilent X090930-2). Cutaneous melanocytes were labeled with anti-gp100 primary antibody (1:100; Abcam ab137078) and Alexa Fluor 555 secondary antibody (4 µg/mL, Thermo Fisher A21428). Nuclei were counterstained with DAPI (1:10,000). Five images were taken for each biological replicate on a Perkin Elmer Vectra automated quantitative pathology imaging system and each image was counted by five blinded reviewers.

### shRNA knock-down

Mission shRNA vectors purchased from Sigma were transiently transfected along with pCMV-VSVG and ps-PAX2 into HEK 293T cells using polyethylenimine (PEI) at a ratio of 3 µL of 10 µg/µL PEI per 1 µg of plasmid (shRNA information provided in **Table S5B**). Viral supernatant was collected 48- and 72-hours post-transfection and filtered through a 0.45 µm syringe filter. Viral supernatant was added to NRAS-null MEFs along with 10 µg/mL polybrene. Fresh media was placed on the cells the following day and 1.5 µg/mL puromycin selection began 48 hours post-infection.

### Adenoviral amplification

RAF NanoBiT constructs were cloned into the pAdTrack shuttle construct. *Pme1*-linearized pAdTrack plasmid was then electroporated into BJ5183-AD-1 cells. The recombined AdEasy vector isolated from the transformed cells was digested with *Pac1* and transfected into HEK 293AD cells using PEI at a ratio of 3 µL of 10 µg/µL PEI per 1 µg of plasmid. Following serial propagation of the virus through HEK 293AD cells, the adenovirus was purified using a CsCl gradient and dialyzed in dialysis buffer (10 mM Tris (pH 8), 2 mM MgCl, 4% sucrose). Purified viral solutions were mixed with glycerol and stored at −80°C.

### NanoBiT assays

Passage four *TN*^*61X*/*X*^ MEFs induced to express NRAS were equally seeded into a 96 well plate. The following day, the cells were placed in DMEM containing 0.5% FBS, 1% penicillin-streptomycin, and 1% glutamine and infected with adenovirus expressing the indicated RAF NanoBiT constructs. The following day, the cells were washed in PBS and placed in DMEM containing 10% FBS, 1% penicillin-streptomycin and 1% glutamine. 48-hours post-infection, the cells were washed with PBS and incubated in serum-free DMEM containing 1% penicillin-streptomycin and 1% glutamine for four hours prior to analysis. Luminescence intensity was assessed using the Nano-Glo Live Cell Assay (Promega N2012). The cells were then fixed in 10% neutral buffered formalin with crystal violet (0.01% w/v) for 30 minutes. Crystal violet-stained plates were imaged on a LI-COR CLx and quantified with Image Studio software. Luminescence intensity was normalized to the crystal violet staining intensity for each well.

### Statistics and reproducibility

Statistical analyses for Kaplan-Meier curves and dot plots were performed using GraphPad Prism version 8.4.3. Survival differences observed in Kaplan-Meier curves were assessed using log-rank (Mantel-Cox) tests. One-way ANOVA was used to analyze dot plots along with a correction for multiple comparisons as stated in each figure legend. Dot plots depict the mean ± s.d. of data acquired from ≥ 3 biological replicates with each dot representing a single replicate. *p* > 0.05 was considered not significant.

## DATA AVAILABILITY

Raw sequencing reads from RNA sequencing are available on NCBI Gene Expression Omnibus with the accession number GSE162124.

## ACKNOWLEDGEMENTS

The authors thank the members of the Genetically Engineered Mouse Modeling Core (GEMMC) at The Ohio State University and the Genomics Services Laboratory (GSL) at Nationwide Children’s Hospital for their technical services. In addition, we would like to thank Emma Crawford, Shannon Gray, Makanko Komara and Suohui Zhang for technical support, Dr. Krista M. D. LaPerle for pathology support, Dr. Dafna Bar-Sagi (NYU) for reagents, and Dr. Martin McMahon (University of Utah) for critical reading of the manuscript. This work was supported by the Damon Runyon Foundation (Innovation Award #38-16 to C.E.B), Pelotonia (B.M.M), and the National Institutes of Health (T32GM068412, F31CA236418 to B.M.M.; P30 CA016058 to the OSUCCC).

## SUPPLEMENTARY FIGURES

**Murphy B**.**M**., **et al**.

**Figure S1:**
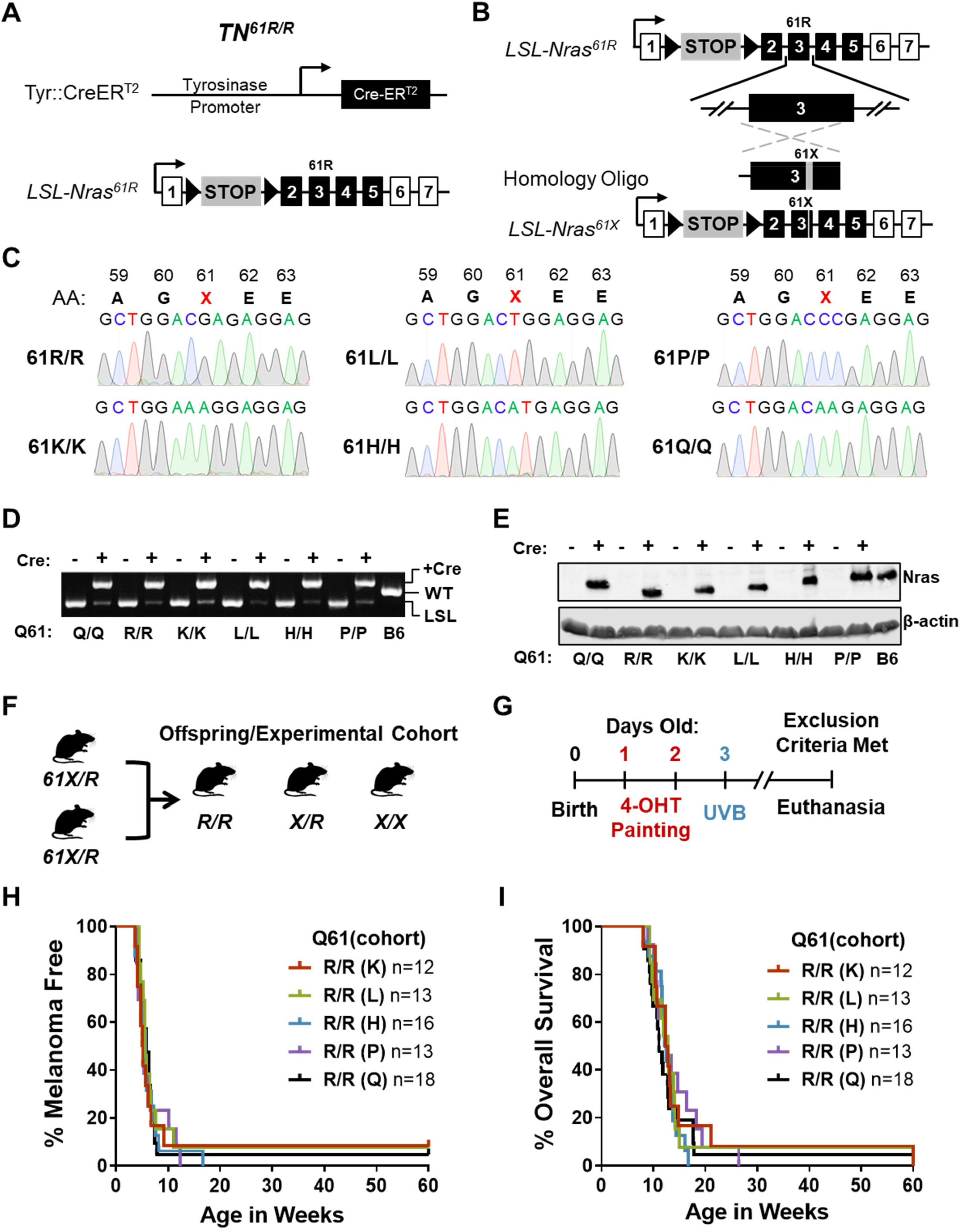
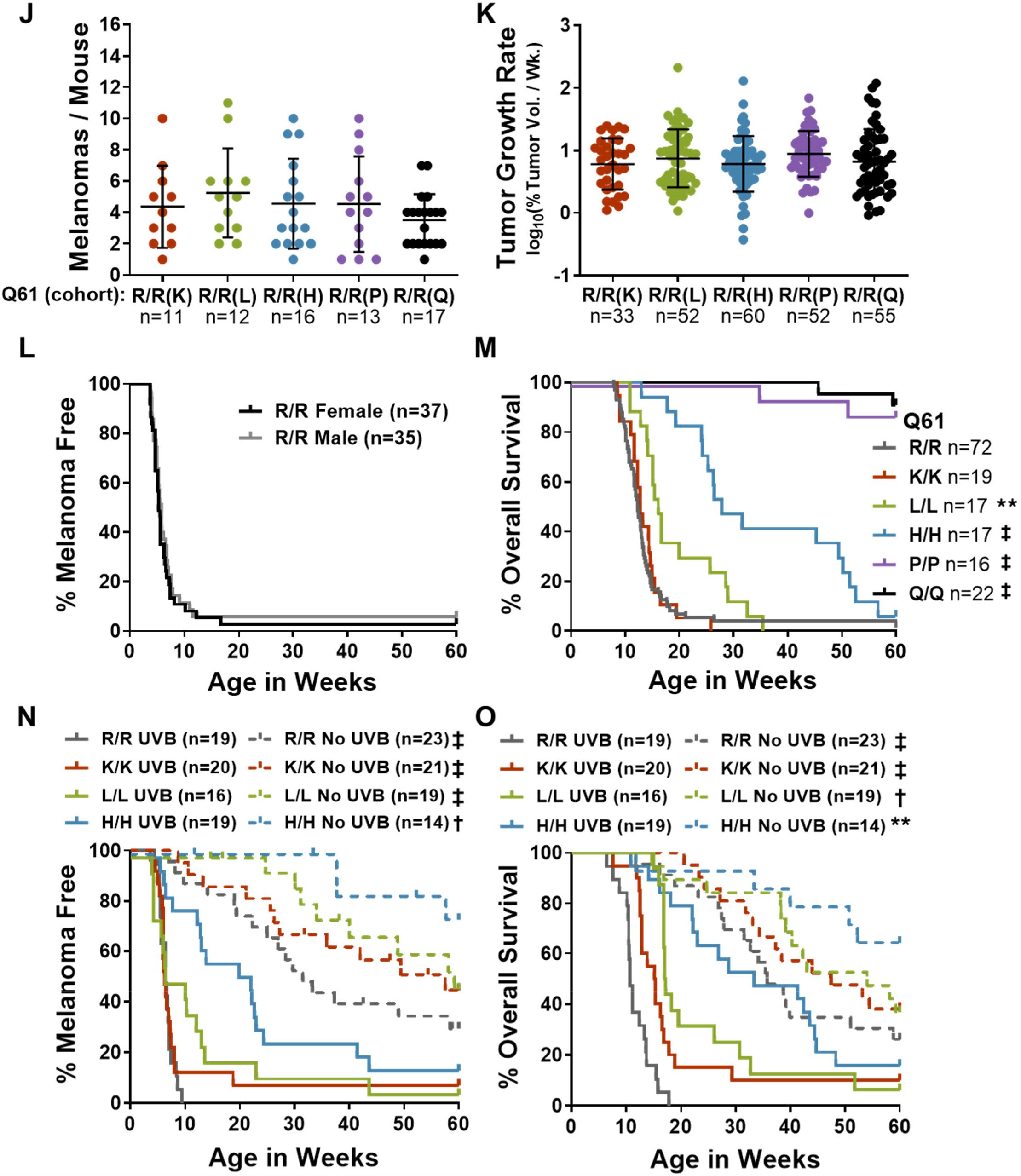
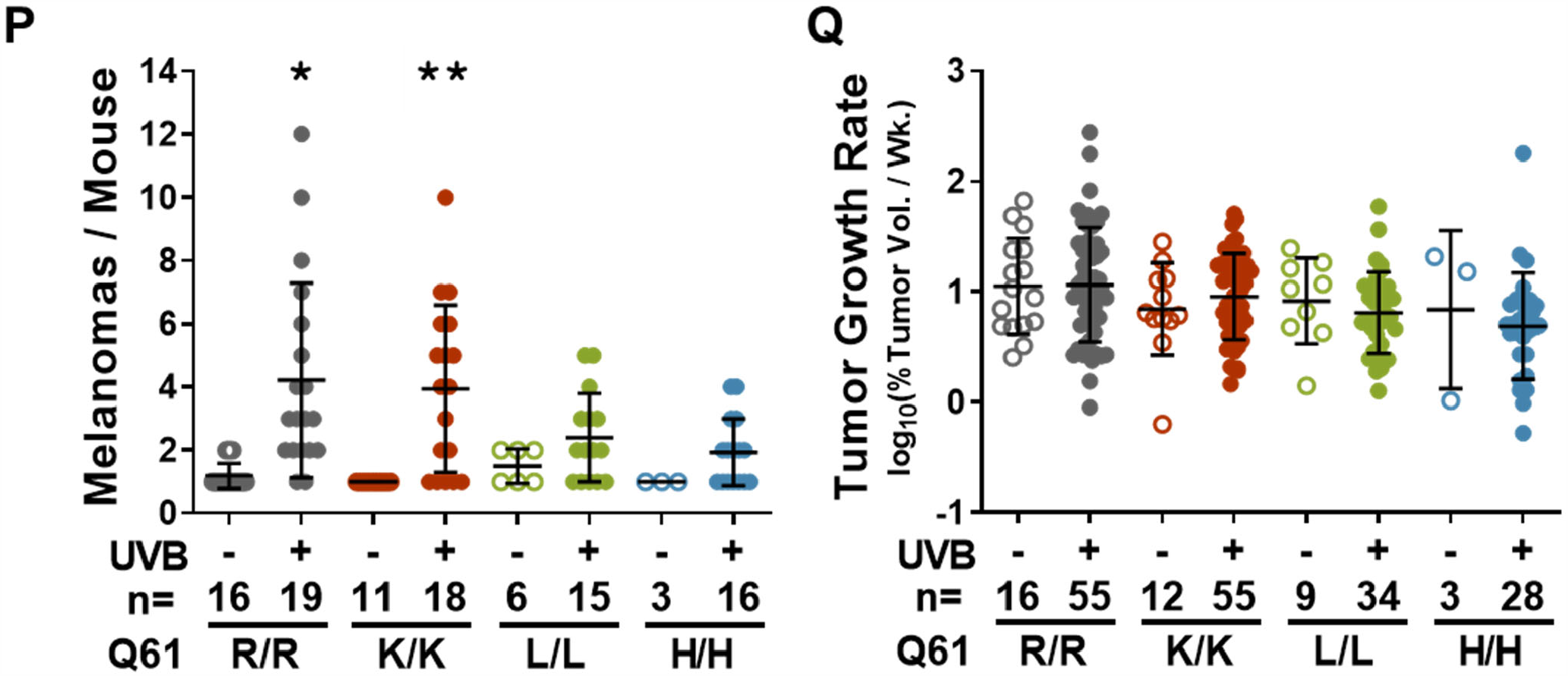
Mouse model generation and experimental design for evaluating the tumorigenicity of NRAS codon 61 mutants. **A**, Diagram of alleles present in the *TN*^*61R*/*R*^model. *TN*^*61R*/*R*^ mice are homozygous for a melanocyte-specific Cre-ER^T2^transgene (Tyr::Cre-ER^T2^) and the *LSL-Nras*^*61R*^ conditional knock-in allele. **B**, Schematic representation of the CRISPR-Cas9-directed strategy used to alter *Nras* codon 61 in *TN*^*61R*/*R*^ embryos. **C**, Sequencing chromatograms of *Nras* exon 3 in MEF DNA isolated from each homozygous *TN*^*61X*/*X*^ model. AA: amino acid. **D**, PCR screen of *LSL* recombination in genomic DNA isolated from murine embryo fibroblasts (MEFs) 72 hours post-infection with adenoviral eGFP (-Cre, control) or Cre (+Cre). **E**, Immunoblot of protein lysates isolated from MEFs 5 days after adenoviral infection as described in ‘D’. **F**, Breeding scheme used to generate experimental litters of *TN*^*61X*/*X*^mice. **G**, Diagram of the experimental treatment protocol. Topical application of 20 mM 4-hydroxytamoxifen (4-OHT) on post-natal days 1 and 2 was used to induce CreER^T2^activity. Ultraviolet-B (UVB) irradiation was administered on post-natal day 3 as a single, 4.5 kJ/m^2^ dose using a fixed position, 16W, 312 nm UVB light source. **H-K**, Melanoma-free survival (H), overall survival (I), total tumor burden (J) and tumor growth rates (K) for *TN*^*61R*/*R*^ mice from each experimental cohort. **L**, Melanoma-free survival of male and female *TN*^*61R*/*R*^ mice. No significant phenotypic differences were detected between cohorts or sexes in log-rank (Mantel-Cox) (H-I, L) and ANOVA analyses with Tukey’s multiple comparison test (J-K). **M**, Overall survival of *TN*^*61X*/*X*^ mice homozygous for the indicated *LSL-Nras* alleles. Log-rank (Mantel-Cox) tests were used to compare the overall survival of each genotype to *TN*^*61R*/*R*^. **N-Q**, Melanoma-free survival (N), overall survival (O), total tumor burden (P) and tumor growth rates (Q) for *TN*^*61X*/*X*^ mice treated with mock or UVB irradiation on post-natal day 3. Log-rank (Mantel-Cox) (N-O) or ANOVA (P-Q) with a Dunnet T3 multiple comparisons test was used to compare measurements between UVB- and mock-treated mice within the same genotype. Adjusted p-values for all comparisons can be found in **Table S1A-B. \*** p< 0.05, **\*\*** p< 0.01, † p< 0.001, ‡ p< 0.0001.

**Figure S2:**
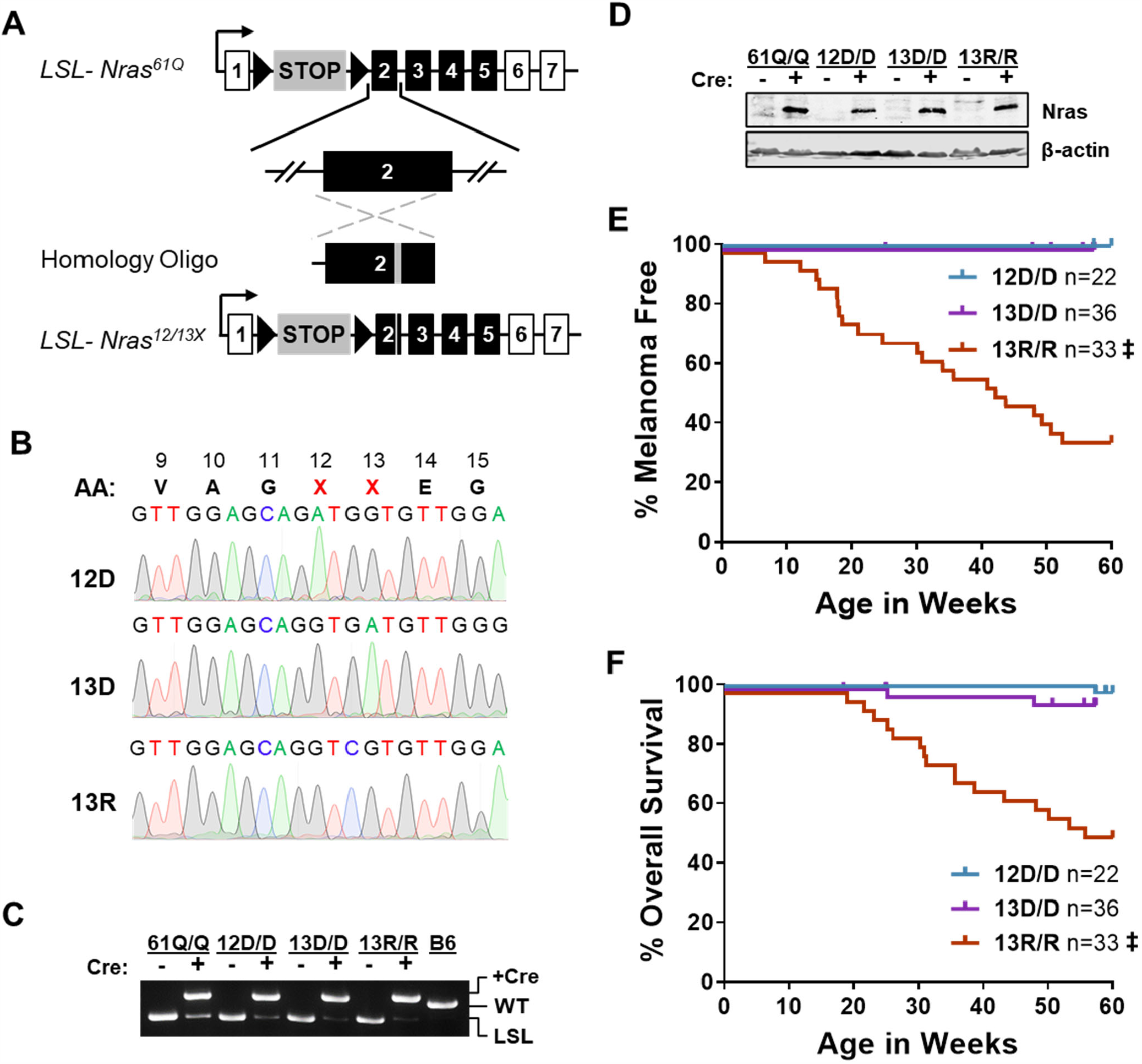
Generation and evaluation of melanoma formation in NRAS codon 12 and 13-mutant mouse models. **A**, Schematic representation of the CRISPR-Cas9-directed mutagenesis strategy used to modify *Nras* codon 12 or 13 in *TN*^*61Q*/*Q*^ embryos. **B**, Sequencing chromatograms of exon 2 in tissue DNA obtained from each homozygous *TN*^*12XX*^or *TN*^*13XX*^ model. A silent G/A mutation at codon 15 occurred during the generation of the *LSL-Nras*^*12D*^ and *LSL-Nras*^*13R*^ alleles. **C-D**, Homozygous *TN*^*12XX*^ or *TN*^*13XX*^ MEFs were infected with adenovirus as described in Figure S1D. PCR shows the recombination of each *LSL-Nras* allele (C) and an immunoblot confirms protein expression (D). **E-F**, Melanoma-free survival (E) and overall survival (F) of mice expressing the indicated melanocyte-specific NRAS^G12^ or NRAS^G13^ mutants. Log-rank (Mantel-Cox) tests were used to compare each genotype to *TN*^*12D*/*D*^. ‡ p< 0.0001.

**Figure S3:**
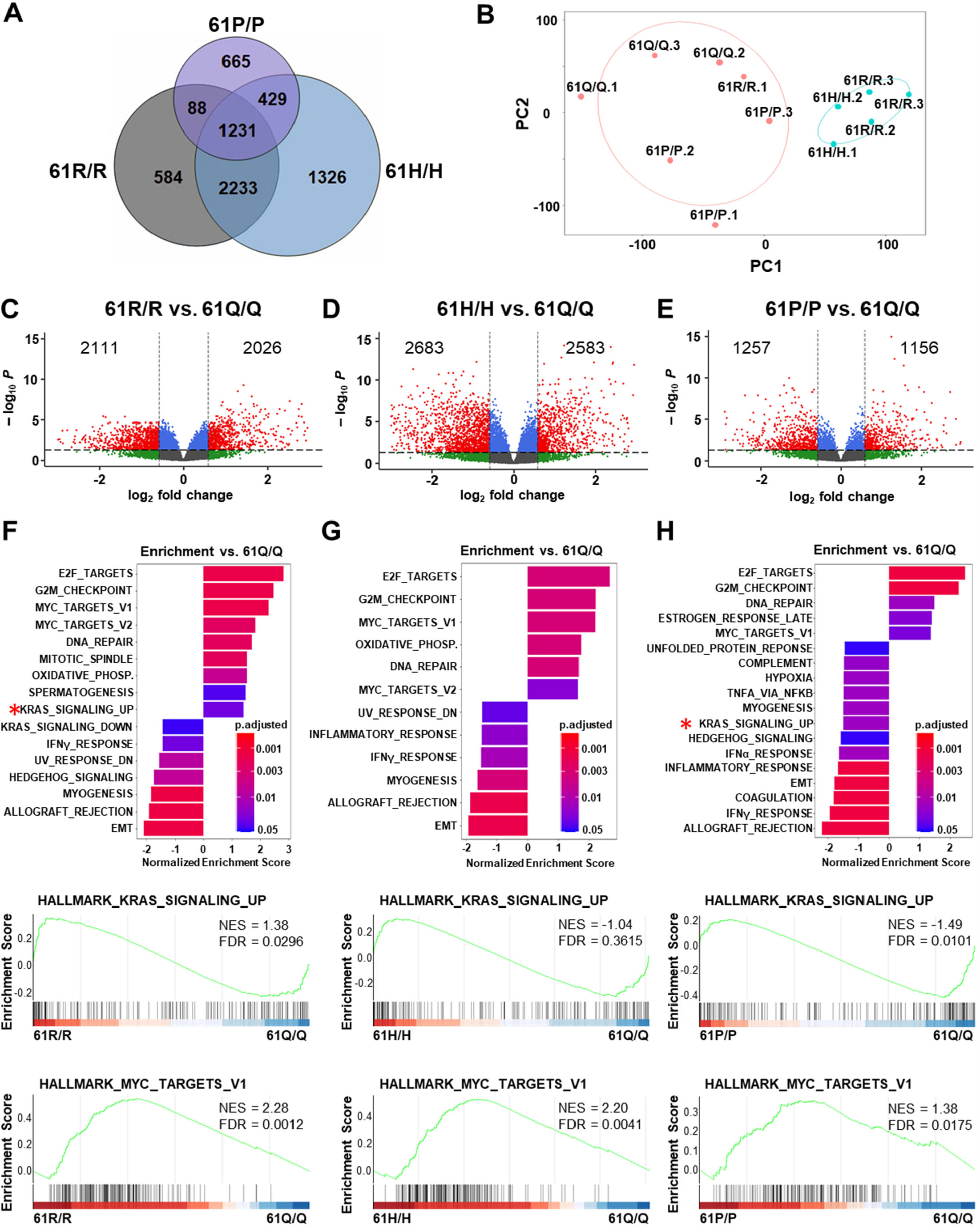
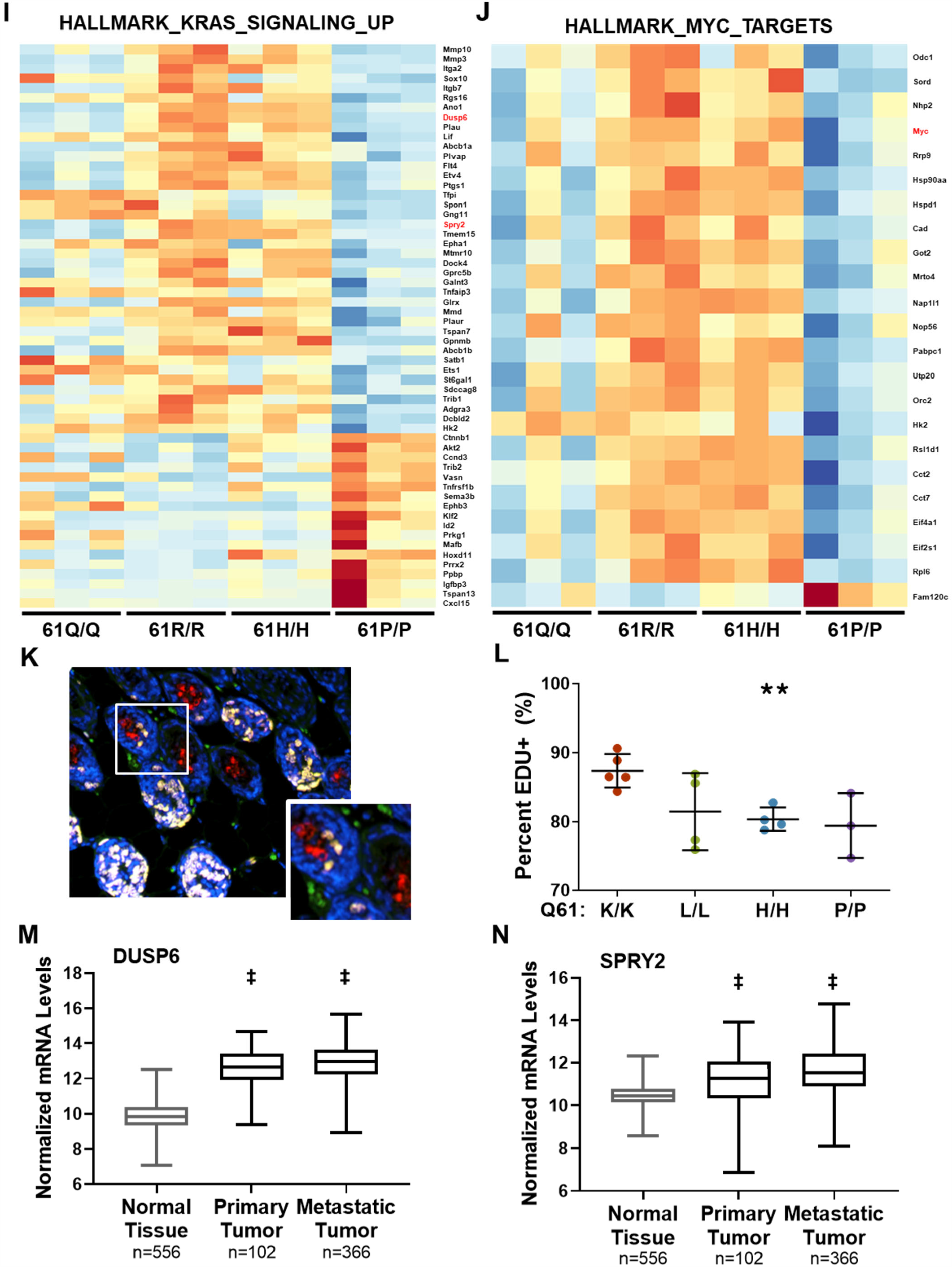
Melanomagenic NRAS mutants drive proliferation and the enrichment of hallmark genes associated with increased KRAS signaling. **A**, Venn diagram depicting the overlap of genes with differential expression in MEFs expressing NRAS^61Q/Q^ and each of the denoted NRAS^61X/X^ mutants. **B**, Principal component analysis of NRAS-mutant MEF samples analyzed via RNA-seq. Samples were separated into two groups based on hierarchical clustering. **C-E**, Volcano plot of differentially expressed genes in NRAS^61R/R^ and NRAS^61Q/Q^MEFs (C), NRAS^61H/H^ and NRAS^61Q/Q^ MEFs (D) or NRAS^61P/P^ and NRAS^61Q/Q^ MEFs (E). **F-H**, (top) Bar plot showing the enrichment of Hallmark gene sets (p-adjusted < 0.05) in MEFs expressing NRAS^61R/R^ versus NRAS^61Q/Q^ (F), NRAS^61H/H^ versus NRAS^61Q/Q^ (G) or NRAS^61P/P^ versus NRAS^61Q/Q^ (H). (bottom) Enrichment plots generated from differentially expressed genes associated with the Hallmark gene sets: KRAS_SIGNALING_UP or MYC_TARGETS_V1. **I-J**, Heatmap of differentially expressed genes from the KRAS_SIGNALING_UP (A) or MYC_TARGETS_V1 (B) gene sets in NRAS^61R/R^ versus NRAS^61P/P^ MEFs (p-adjusted < 0.05). **K**, Representative image of EdU (proliferation, green) and gp100 (melanocyte, red) co-staining in skin harvested from a ten-day old mouse. **L**, Dot plot showing percent EdU positivity in melanocytes from 10-day old *TN*^*61X*/*X*^ mouse skin. Each dot represents one biological replicate. ANOVA with a Tukey post-test was used to compare data from *Nras*^*61K*/*K*^ and *Nras*^*61X*/*X*^ cells. **M-N**, Box plots of *DUSP6* (M) and *SPRY2* (N) human gene expression in normal skin tissue (GTEX) versus primary and metastatic cutaneous melanoma (TCGA). Data were obtained from the UCSC Xena platform (https://doi.org/10.1038/s41587-020-0546-8). ANOVA with a Tukey post-test was used to compare expression levels from normal tissue to primary and metastatic tumors. **\*\*** p< 0.01, ‡ p< 0.0001.

**Figure S4:**
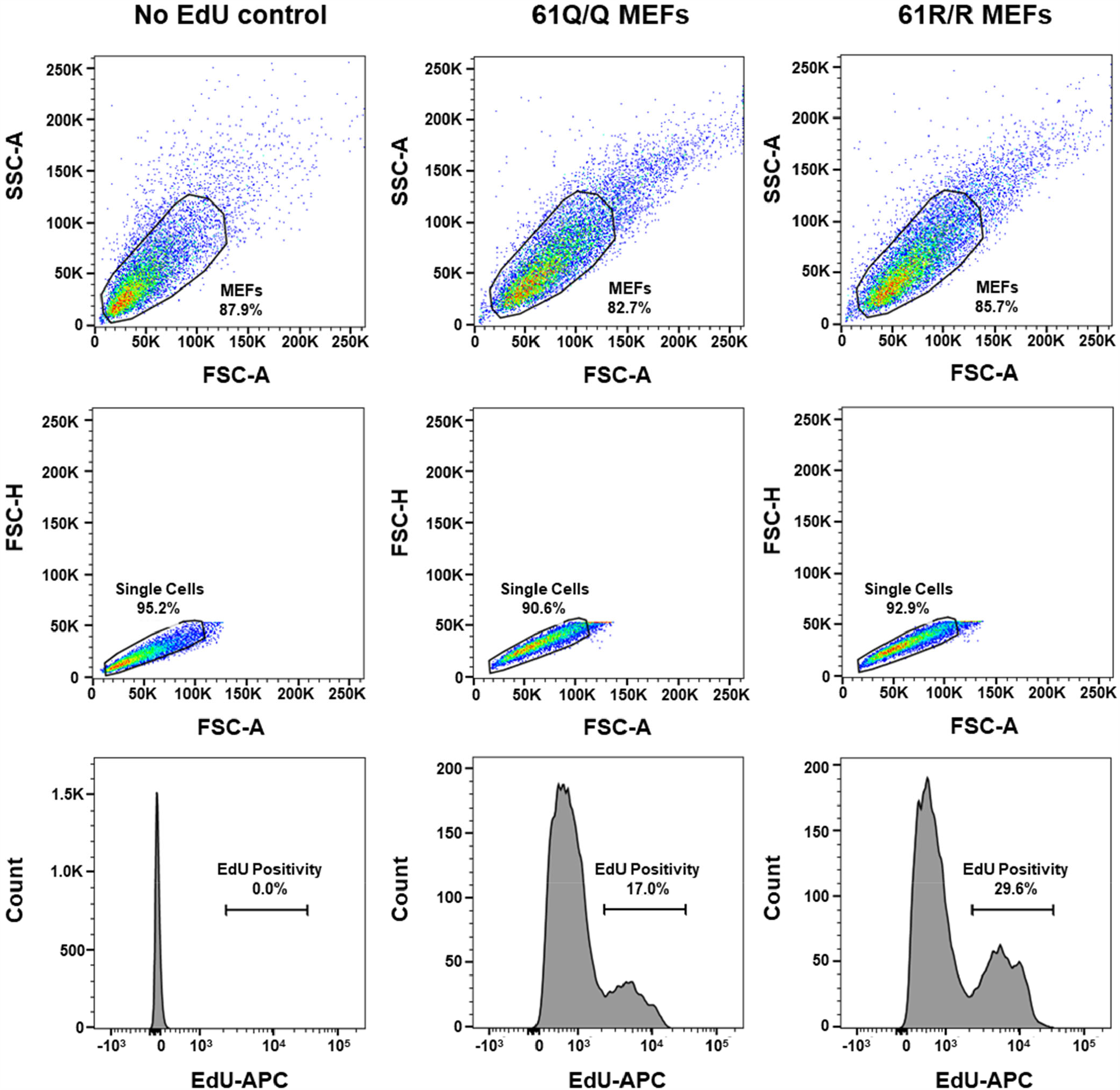
Gating strategy for flow cytometric analysis of *in vitro* EdU incorporation in MEFs. MEFs, labeled with EdU conjugated to Alexa Fluor 555, were analyzed on a BD LSR Fortessa flow cytometer. The initial population of MEFs was selected by gating based on FSC-A by SSC-A (top). Cell doublets were removed by gating for single cells in a FSC-H by FSC-A plot (middle). Finally, a histogram of counts by APC-A intensity was used to determine the percent of EdU positive cells in each population of MEFs (bottom).

